# Biomarker Discovery via Integrative Multi-omics for Children exposed to Humidifier Disinfectant

**DOI:** 10.1101/2025.09.04.674118

**Authors:** Jaeho Ji, Ahrum Son, Mi-Jin Kang, Jeonghun Yeom, Hyun Ju Yoo, Kwoneel Kim, Jeong-Hyun Kim, Hea Young Oh, Shin Ah Kim, So-Yeon Lee, Seung-Hwa Lee, Soo-Jong Hong, Hyunsoo Kim

**Affiliations:** Graduate School of Life Sciences, College of Bioscience and Biotechnology, Chungnam National University, Daejeon, Republic of Korea; Graduate School of Medical Science, Department of Medical Science Convergence, University of Ulsan, Ulsan, Republic of Korea; Humidifier Disinfectant Health Center, Asan Medical Center, Seoul, Republic of Korea; Prometabio Research Institute, Prometabio Co., Ltd, Gyeonggi-do, Republic of Korea; Department of Convergence Medicine and Department of Digital Medicine, Asan Institute for Life Sciences, Asan Medical Center, University of Ulsan College of Medicine, Seoul, Republic of Korea; Department of Biology, Kyung Hee University, Seoul, Republic of Korea; Department of Medicine, University of Ulsan College of Medicine, Seoul, Republic of Korea; PHI Digital Healthcare, Seoul, Republic of Korea; Institute for Innovation in Digital Healthcare, Yonsei University, Seoul, Republic of Korea; Department of Pediatrics, Humidifier Disinfectant Health Center, National Medical Center, Seoul, Republic of Korea; Department of Convergent Bioscience and Informatics, College of Bioscience and Biotechnology, Chungnam National University, Daejeon, Republic of Korea; Department of Bio-AI Convergence, Chungnam National University, Daejeon, Republic of Korea; Institute of Biotechnology, Chungnam National University, Daejeon, Republic of Korea; Protein Design Institute, Chungnam National University, Daejeon, Republic of Korea; SCICS, Daejeon, Republic of Korea

**Keywords:** Humidifier disinfectant, Multi-omics, Multi-marker panel, Integrin-related signaling, Chronic respiratory conditions

## Abstract

**Rationale:** Exposure to humidifier disinfectants has been linked to an array of pulmonary disorders and diminished lung functionality particularly reduced Forced Vital Capacity (FVC).

**Objectives:** This investigation sought to identify diagnostic biomarkers for early detection of children at elevated risk of developing chronic respiratory conditions following such exposure.

**Methods:** Our research employed a comprehensive multi-omics strategy analyzing 70 pediatric patients alongside 10 controls, seamlessly integrating clinical assessments with transcriptomics, methylomics, proteomics, and metabolomics data. The analytical framework utilized a sophisticated combination of Non-negative Matrix Factorization (NMF), Multi-Omics Factor Analysis (MOFA), and advanced machine learning algorithms.

**Measurements and Main Results:** NMF clustering uncovered distinctive protein expression patterns associated with integrin-mediated signaling pathways and immune response mechanisms. Complementarily, MOFA identified latent factors correlating with lung function metrics, highlighting critical molecular pathways involved in integrin cell surface interactions and lipid metabolism regulation. Machine learning-based analysis facilitated the development of a multi-marker panel–comprising IGHV2-70, LysoPC (16:0), and hexadecyl ferulate–which achieved 81.46% accuracy in identifying pulmonary dysfunction cohort.

**Conclusions:** These findings suggest that alterations in integrin-related signaling networks and dysregulation of lipid metabolism play pivotal roles in mediating the long-term pulmonary consequences of humidifier disinfectant exposure. The proposed multi-marker panel offers significant potential for enhanced risk stratification and timely therapeutic intervention.

**At a Glance Commentary:** *Scientific Knowledge on the Subject:* Extensive epidemiological evidence has established the causal relationship between humidifier disinfectant exposure and pulmonary dysfunction; however, clinically validated biomarkers for predicting chronic lung disease progression remain limited. Pediatric populations demonstrate unique pathophysiological mechanisms distinct from adults, highlighting the critical necessity for biomarker identification grounded in comprehensive molecular understanding. Despite advances in omics technologies, recent investigations have encountered significant obstacles in achieving deeper mechanistic insights, predominantly attributable to methodological constraints in harmonizing clinical phenotypes with high-dimensional molecular datasets.

*What This Study Adds to the Field:* This investigation elucidates the fundamental contributions of integrin-mediated signaling cascades and lipid metabolic networks to persistent pulmonary dysfunction following humidifier disinfectant exposure. Our analyses revealed coordinated regulation of integrin signaling pathways and immune response networks through NMF clustering, indicating dynamic temporal evolution of inflammatory responses during chronic disease progression, with temporally distinct molecular signatures identified across discrete observation intervals. Multi-omics factor analysis (MOFA) corroborated integrin pathway dysregulation while additionally uncovering systematic suppression of lipid metabolic processes. Furthermore, machine learning algorithms enabled development of a robust three-component biomarker panel—encompassing IGHV2-70, LysoPC (16:0), and hexadecyl ferulate—demonstrating 81.46% classification accuracy for pulmonary dysfunction phenotypes. Collectively, these findings substantially advance mechanistic understanding of chronic lung injury in vulnerable pediatric cohorts and identify clinically relevant biomarkers with translational potential for risk stratification and therapeutic targeting in clinical practice.

## Introduction

The 2006 emergence of an unidentified respiratory disease in Korea heralded a significant public health crisis that would later be traced to a specific source: humidifier disinfectants (HDs). Following the Korean Centers for Disease Control and Prevention (KCDC) 2011 ban on these products, new cases declined precipitously (1). Nevertheless, the long-term consequences of exposure persist, particularly among young children who constitute the majority of survivors (2). Subsequent investigations several chemical constituents in these disinfectants, including polyhexamethylene guanidine (PHMG), oligo (2-(2-ethoxy)ethoxyethyl) guanidinium chloride (PGH), chloromethylisothiazolinone (CMIT), and methylisothiazolinone (MIT). These compounds have been implicated in the development of Humidifier Disinfectant Lung Injury (HDLI) (3). Among these, PHMG has used commonly and emerged as a primary etiological factor in various pulmonary pathologies (4), including inflammation, lung fibrosis (5), and interstitial lung diseases (ILDs) (6). Characterized by pulmonary fibrosis resulting from toxic chemical inhalation (7), HDLI presents histologically with a distinctive bronchiolocentric distribution (8) and clinically manifests as diminished lung function, as measured through Pulmonary Function Testing (PFT) (9). The causal relationship between these disinfectants and the observed pathologies has been substantiated through comprehensive studies conducted on Korean patient cohorts (9, 10). Recent research has expanded the spectrum of associated respiratory conditions to encompass Chronic Obstructive Pulmonary Disease (COPD), asthma, and ILDs (2, 7).

In contrast to their adult counterparts, children exhibit heightened vulnerability to the development of pulmonary pathologies, most notably childhood ILD, following exposure to these disinfectant compounds (1, 11). Compelling evidence from longitudinal studies has revealed that such exposures during critical developmental windows in early childhood precipitate persistent alterations in respiratory function that extend well into later life stage (12). Nonetheless, comprehensive investigation employing multi-omics approaches to elucidate the underlying mechanisms remains at a nascent stage, with significant knowledge gaps yet to be addressed. Preclinical research utilizing rodent models has successfully integrated tripartite omics datasets to identify potential protein and metabolite biomarkers associated with respiratory toxicity induced by PHMG-phosphate, a principal constituent of the implicated disinfectants (13). In the clinical domain, previous comparative analyses of adult and pediatric cohorts have begun to unveil distinctive biological mechanisms of lung pathogenesis that underpin HDLI across these age demographics. However, these investigations remain limited to tissue-based approaches with incomplete multi-omics coverage, whereas our study pioneers the systematic identification of blood-based biomarkers through comprehensive four-layer omics integration (14).

Our study addresses this research lacuna through integrative analysis, employing methodologies at the forefront of multi-layer omics research–encompassing transcriptomics, methylomics, proteomics, and metabolomics–and advanced bioinformatics approaches. These include non-negative matrix factorization (NMF), multi-omics factor analysis (MOFA), and advanced machine learning techniques. NMF functions as a dimensionality reduction method that extracts key features while maintaining non-negativity constraints (15). Originally developed for computational applications such as text mining and image processing, it has been adapted for bioinformatics applications, including gene expression clustering and cancer mutation analysis (16). MOFA provides a framework for identifying factors that elucidate variation in multi-omics datasets through a Bayesian probabilistic model for dimensionality reduction (17, 18). Complementing these approaches, machine learning methodologies have emerged as particularly valuable in the bioinformatics domain for biomarker panel classification, effectively transcending the inherent limitations associated with single biomarker (19, 20).

Currently, medical support for individuals exposed to humidifier disinfectants primarily focuses on HDLI. However, pediatric patients with exposure history face an elevated risk of developing chronic lung diseases in adolescence and adulthood due to early pulmonary damage. This underscores the necessity for additional medical support targeting a broader spectrum of chronic respiratory conditions. The primary objective of our study is to identify biomarkers that could facilitate the early diagnosis of patients at high risk for developing these respiratory conditions. This research aims to contribute to the growing body of knowledge surrounding the long-term health impacts of humidifier disinfectant exposure, with a particular focus on pediatric populations. By leveraging advanced multi-omics and bioinformatics techniques, we seek to uncover novel biomarkers and pathways that may inform future diagnostic and therapeutic strategies for this vulnerable patient group.

## Methods

### Study Population and Design

We examined 70 pediatric patients (aged <18 years) enrolled in the Humidifier Disinfectant Health Monitoring Program I, with confirmed exposure to HDs containing predominantly (> 50%) PHMG or PGH, who underwent serial pulmonary function testing (≥4 assessments) during longitudinal follow-up. Based on pulmonary function test results, participants were stratified into four distinct groups: Severe group (severe lung injury, n=10) designated as critical lung injury, exhibited decreased lung function such as FVC and FEV_1_ accompanied by notable symptoms; Restrictive group (restrictive lung disease, n=19) presented with attenuated symptoms but similar patterns of diminished lung function such as FVC; Obstructive group (obstructive lung disease, n=21) demonstrated high risk for developing COPD or asthma in adulthood with decreased FEV_1_; and Normal group (normal lung function, n=20) maintained normal lung function despite humidifier disinfectant exposure. A Control group (control cohort, n=10) comprising individuals with neither humidifier disinfectant exposure nor diagnosed respiratory disease was included for comparative analysis. These participants were recruited from the Cohort for Childhood Origin of Asthma and Allergic Diseases (COCOA), a prospective national birth cohort study following healthy Korean newborns from birth (21, 22). Restrictive group and obstructive group were designated as primary target groups due to their elevated likelihood of developing chronic lung disease in adulthood. Although Severe group exhibited similar potential, these patients were excluded from the target classification given their pre-existing severe lung damage. Longitudinal assessment was conducted at two distinct time points during 4-5 years monitoring period: ‘First’ (baseline pulmonary function assessment), and ‘Last’ (final follow-up visit). Multi-omics data obtained from each patient at both time points enabled comprehensive evaluation of disease progression and potential biomarkers. This methodology facilitated a nuanced understanding of long-term effects of humidifier disinfectant exposure on pulmonary function, while providing molecular-level insights into underlying pathophysiological mechanisms. This study obtained approval from the Institutional Review Board (IRB) of Asan Medical Center (2018-1406), and informed consent was waived by the IRB.

### Spirometry

Pulmonary function tests were performed using a calibrated spirometer (Vmax229D; SensorMedics) at the Humidifier Disinfectant Health Center of Asan Medical Center, Seoul. All procedures strictly followed the standardized guidelines established by the American Thoracic Society (ATS) and European Respiratory Society (ERS) in 2019 (23). Several key respiratory parameters were assessed, including Forced Expiratory Volume in one second (FEV₁), which measures the volume of air forcefully expelled during the first second of expiration. Forced Vital Capacity (FVC), was also evaluated, representing the total air volume expelled during a complete forced expiratory maneuver. The FEV₁/FVC ratio was calculated to assess the relationship between airflow limitation and lung volume. Additionally, Forced Expiratory Flow over the middle half of the FVC (FEF_25-75%_) was measured, which reflects average flow rates across the middle 50% of the FVC and provides valuable insights into small airway function. These spirometric parameters served as the foundation for clinical classification of patient groups according to established PFT interpretation criteria (24) and constituted essential clinical variables in the subsequent integrative analysis, enabling comprehensive assessment of pulmonary function across the study cohort.

### Multi-omics Data Collection

Blood samples from study participants were systematically processed to generate a comprehensive multi-omics dataset. For proteome analysis, plasma samples underwent S-trap digestion followed by Sequential Window Acquisition of All Theoretical Mass Spectra (SWATH-MS) analysis. A ZenoTOF 7600 mass spectrometer (SCIEX) coupled with a Vanquish Neo UHLPC system (Thermo Fisher Scientific) was employed for all measurements. Samples were analyzed using Zeno SWATH, a data-independent acquisition (DIA) technique that enhances sensitivity through Zeno trap pulsing via a linear ion trap, as previously described (25). Data processing was conducted using SwissProt 2022.04 and DIA-NN v1.8.1. While 999 proteins were initially identified, after excluding seven proteins lacking annotations and those with excessive missing values (detailed in the Data Preprocessing section), 822 proteins were retained for subsequent analyses. For metabolome characterization, non-targeted profiling was performed using an Ultimate 3000/LTQ-Orbitrap XL system. Metabolic features were identified through comparative analysis against Metlin, Kyoto Encyclopedia of Genes and Genomes (KEGG), and Human Metabolome Database (HMDB), utilizing the in-house database mzCloud. Features were classified according to their metabolic annotation level as Tentative Structure and/or Putative Identification (level 2-3) (26), resulting in 752 distinct metabolic features. Transcriptome data from the peripheral blood preserved in PAXgene tubes were generated using the GeneChip® Human Gene 2.0 ST microarray technology, with availability restricted to the late data collection phase. Methylome profiling was conducted using the Illumina Infinium MethylationEPIC v2.0 BeadChip platform in conjunction with a BeadXpress reader, ensuring high-throughput and precise epigenetic characterization. This integrated multi-omics approach enabled a comprehensive examination of the molecular landscape associated with humidifier disinfectant exposure and its potential long-term effects on pulmonary function.

### Data Preprocessing

A standardized protocol was implemented for preprocessing all omics datasets to ensure consistency and reliability. Initially, log_2_ transformation was applied to all omics data to achieve variance stabilization without additional normalization procedures. For the proteome dataset, which contained missing values (NAs), a rigorous filtering strategy was employed. Features with 30 or more NAs were excluded from subsequent analyses to maintain data integrity. The remaining missing values were imputed using the K-Nearest Neighbors (KNN) method, a robust approach that preserves the underlying data structure while minimizing potential biases introduced by incomplete data. In parallel, clinical parameters derived from pulmonary function tests (PFTs) were subjected to Z-score normalization, standardizing the scale of diverse variables to facilitate meaningful cross-measurement comparisons. This comprehensive preprocessing framework was designed to optimize data quality and prepare the multi-omics datasets for integrative analyses, thereby enhancing the reliability and interpretability of findings related to humidifier disinfectant exposure and its impact on pulmonary function.

### Clustering Analysis

A comprehensive clustering analysis was performed on multi-omics data from 70 subjects at each time point using the scikit-learn library in Python (27). Initially, K-means clustering was employed with the optimal number of clusters determined through the elbow method, which evaluates cluster quality based on average and median centroid data (28). To examine hierarchical relationships within the data, dendrograms were constructed using Ward linkage, while principal component analysis (PCA) was utilized to visualize overall data distribution. Non-negative Matrix Factorization (NMF)-based clustering was then applied to specific multi-omics datasets: proteome and metabolome data for the early time point (First), and proteome, metabolome, and transcriptome data for the late time point (Last). The extensive methylome dataset was excluded from this analysis due to computational constraints. For NMF clustering, the NMF library in R from CRAN (29) was utilized, with appropriate processing of negative values in the log_2_-transformed data. The Brunet method (30) was implemented with 50 runs (nrun) and rank values ranging from 2 to 10. The optimal number of clusters was determined using the cophenetic correlation coefficient, which measures clustering stability. Finally, data distribution within the identified clusters was verified through consensus map visualization. This multi-faceted clustering approach enabled robust exploration of the underlying structure within the multi-omics datasets, revealing important patterns and relationships associated with humidifier disinfectant exposure and its long-term effects on pulmonary function.

### Differential Expression Proteins and Pathway Enrichment Analysis

Following the identification of clusters through NMF-based clustering, the distribution of clinical indicators within each cluster was meticulously examined to establish appropriate comparison groups. Differential expression analysis was subsequently performed using the DEP library in R (31). For data normalization, Variance Stabilizing Normalization (VSN) was applied to the log_2_-transformed preprocessed data. The differentially expressed protein (DEP) analysis employed rigorous criteria, including a log_2_ fold change threshold of 1.5 and a Benjamini-Hochberg False Discovery Rate (FDR) alpha of 0.05. DEPs were classified as up- regulated or down-regulated based on their cluster ratio. To interpret the biological significance of identified DEPs, functional enrichment analysis was conducted using g:Profiler, a comprehensive web-based toolset (32). This enrichment analysis incorporated queries across multiple databases, including Gene Ontology (GO), Kyoto Encyclopedia of Genes and Genomes (KEGG), and Reactome. To address multiple testing corrections the Benjamini-Hochberg FDR was consistently applied with an alpha threshold of 0.05 across all database queries. This systematic analytical approach enabled identification of significantly altered protein expression patterns and their associated biological pathways, revealing key molecular mechanisms underlying the long-term effects of humidifier disinfectant exposure on pulmonary function.

### Multi-Omics Factor Analysis (MOFA)

Multi-Omics Factor Analysis (MOFA) was conducted on the comprehensive dataset at each time point to identify latent factors driving variability across different omics layers. Given the computational demands of the extensive methylome data, a feature selection approach was implemented, filtering methylome features based on a fold change threshold of >1.5 or <0.66, which yield a refined set of 1,475 features for analysis. To facilitate nuanced understanding of humidifier disinfectant exposure effects, three comparison groups were established: Severe (severe lung injury), Target (encompassing restrictive and obstructive lung disease), Normal (normal lung function). This stratification enabled identification of molecular characteristics common to humidifier disinfectant exposure as well as those differentiating between clinical presentations (17, 18).The MOFA model was constructed using 15 factors with default parameters through the MOFA2 package v1.14.0 in R/Bioconductor. Factors that contributed no variance to the model were subsequently excluded from downstream analyses. Using the MOFA framework, correlations between each latent factor and clinical data were identified, providing insights into relationships between molecular signatures and observed clinical outcomes. For factors specifically related to lung function, pathway enrichment analysis was conducted using the built-in library within the MOFA framework. This integrative analytical approach allowed comprehensive exploration of the complex interplay between multi-omics data and clinical manifestations in the context of humidifier disinfectant exposure, revealing novel biological mechanisms and potential therapeutic targets.

### Machine Learning Analysis

A comprehensive machine learning analysis was conducted using RapidMiner Studio v9.10.010, encompassing data preprocessing, model construction, and training phases. The study employed an extensive array of 18 distinct machine learning algorithms, including Deep Learning (DL), Linear Discriminant Analysis (LDA), AutoMLP (MLP), Decision Stump (DS), Decision Tree (DT), Generalized Linear Model (GLM), Gradient Boosted Trees (GBT), K-Nearest Neighbor (KNN), Naïve Bayes (NB), Naïve Bayes Kernel (kNB), Neural Net (NN), Quadratic Discriminant Analysis (QDA), Random Forest (RF), Random Tree (RT), Regularized Discriminant Analysis (RDA), Rule Induction (RI), Single Rule Induction (SRI), and Support Vector Machine (SVM). The training dataset comprised omics data from patients’ early time points, subjected to the same filtering criteria applied in the Multi-Omics Factor Analysis (MOFA). Model evaluation was conducted using a 3-fold cross-validation approach, implemented through a forward selection method. Performance assessment encompassed various metrics, with a particular emphasis on accuracy. In accordance with the MOFA protocol and to ensure methodological consistency, participants were stratified into three comparison groups based on identical classification criteria: Severe, Target, and Normal categories. This stratification aimed to identify features capable of effectively discriminating between these clinically relevant groups. This multi-model machine learning approach facilitates a robust exploration of the complex relationships within the multi-omics dataset, potentially revealing novel biomarkers and predictive features associated with different outcomes following humidifier disinfectant exposure.

### Statistical Analysis

To validate the temporal associations of markers identified through Machine Learning analyses between the initial (First) and final (Last) data points, a series of statistical tests was conducted. Initially, the Shapiro-Wilk normality test was employed using the Stats library in R v4.4.1 to assess the normality of data distribution for each marker. Subsequently, statistical tests were conducted to assess the equality of variance between the initial (First) and final (Last) data distributions. Bartlett’s test was applied to features exhibiting a normal distribution (met294), while the Brown-Forsythe test was employed for features with non-normal distributions in R v4.4.1. For all features deviating from normality by failing to meet one or more conditions, the non-parametric Wilcoxon signed-rank test was performed using GraphPad v10.0. This approach ensured robust statistical inference across diverse data distributions. Furthermore, correlation analysis was conducted to evaluate the associations between all markers and clinical variables. Spearman’s rank correlation test was performed to examine the relationships between markers and clinical variables for lung function (FEV_1_, FVC, FEV_1_/FVC, and FEF_25-75%_), with all variables normalized using z-score normalization. In the context of Non-negative Matrix Factorization (NMF) and Multi-Omics Factor Analysis (MOFA), statistical analyses were conducted using the algorithms inherent to their respective built-in libraries. This integrated statistical approach allows for a comprehensive evaluation of the temporal stability and significance of identified markers, enhancing the reliability of findings in the context of long-term effects of humidifier disinfectant exposure on pulmonary function.

### Network Analysis

Initially, STRING v12.0 (33) (https://string-db.org/) was employed to assess the overall interaction network of the identified biomarkers. The full STRING network was visualized for the remaining 12 biomarkers, excluding two metabolites and three proteins that were not mapped. The network was configured with a minimum interaction score to high confidence (0.7), and three additional nodes were introduced. Subsequently, pathway enrichment analysis was conducted on the entire human Reactome pathway for 15 protein biomarkers that were converted to gene names using the ReactomeFIViz app v8.0.9. Metabolomic biomarkers were excluded due to their absence in the Reactome database. Functional interaction (FI) networks were obtained for the top three pathway categories based on FDR values. Among these, platelet-related pathways were excluded due to the absence of FI networks. For network filtering and effective visualization, each FI network was reconstructed by inputting its nodes into stringApp (34) v2.2.0’s protein query. A confidence threshold of 0.7 was applied and the physical subnetworks were presented to enhance interpretability. In plasma lipoprotein-related pathways, only two clusters containing protein markers were identified using Markov clustering (MCL) (35) to simplify the representation of complex network relationships. All pathway network analyses were performed in Cytoscape v3.10.3.

## Result

### Longitudinal Study and Analytical Workflow

To investigate the clinical characteristics of patients exposed to humidifier disinfectants, we conducted an integrative analysis utilizing multi-omics and clinical data from individuals enrolled in a longitudinal monitoring cohort. Demographic and clinical characteristics of the 80 enrolled children are presented in Table 1. The study cohort had a mean age of 9 years with equal sex distribution (50.0%, female). Among the 70 participants diagnosed with HDLI, mean HD exposure concentration was 126.02 μg/m³, with a cumulative exposure duration of 5002.86 hours (Table 1). Our analytical workflow was structured into three sequential phases (Figure 1A). First, patients were stratified into distinct groups based on pulmonary function parameters. Second, biomarker discovery was performed through comprehensive bioinformatics analysis integrating multi-omics and clinical datasets, employing three complementary frameworks: NMF-based clustering, Multi-Omics Factor Analysis (MOFA), and machine learning methodologies. Third, identified biomarker candidates were validated for clinical relevance and network-level associations. Paired longitudinal data were collected for each patient at two time points: initial presentation (“First”) and follow-up assessment (“Last”). Based on clinical criteria and analytical constraints, three comparison groups were established: Severe (patients with severe pulmonary dysfunction), Target (patients with restrictive and obstructive patterns), and Normal (control group with normal pulmonary function) (Figure 1B). This three-group classification was specifically designed to accommodate the technical limitations of MOFA+, which supports a maximum of three groups in multi-group analysis mode (18). For NMF-based clustering analysis, the comparison framework was adapted to a binary classification system (High-risk versus Control) due to methodological constraints of the clustering approach. we implemented a comprehensive analytical strategy that integrated multi-layered datasets to identify clinically relevant biomarkers and functional network interactions (Figure 1C). This integrative approach was designed to capture the multi-faceted pathophysiological characteristics associated with humidifier disinfectant exposure, providing mechanistic insights through the convergence of multiple data modalities.

**Figure 1.**
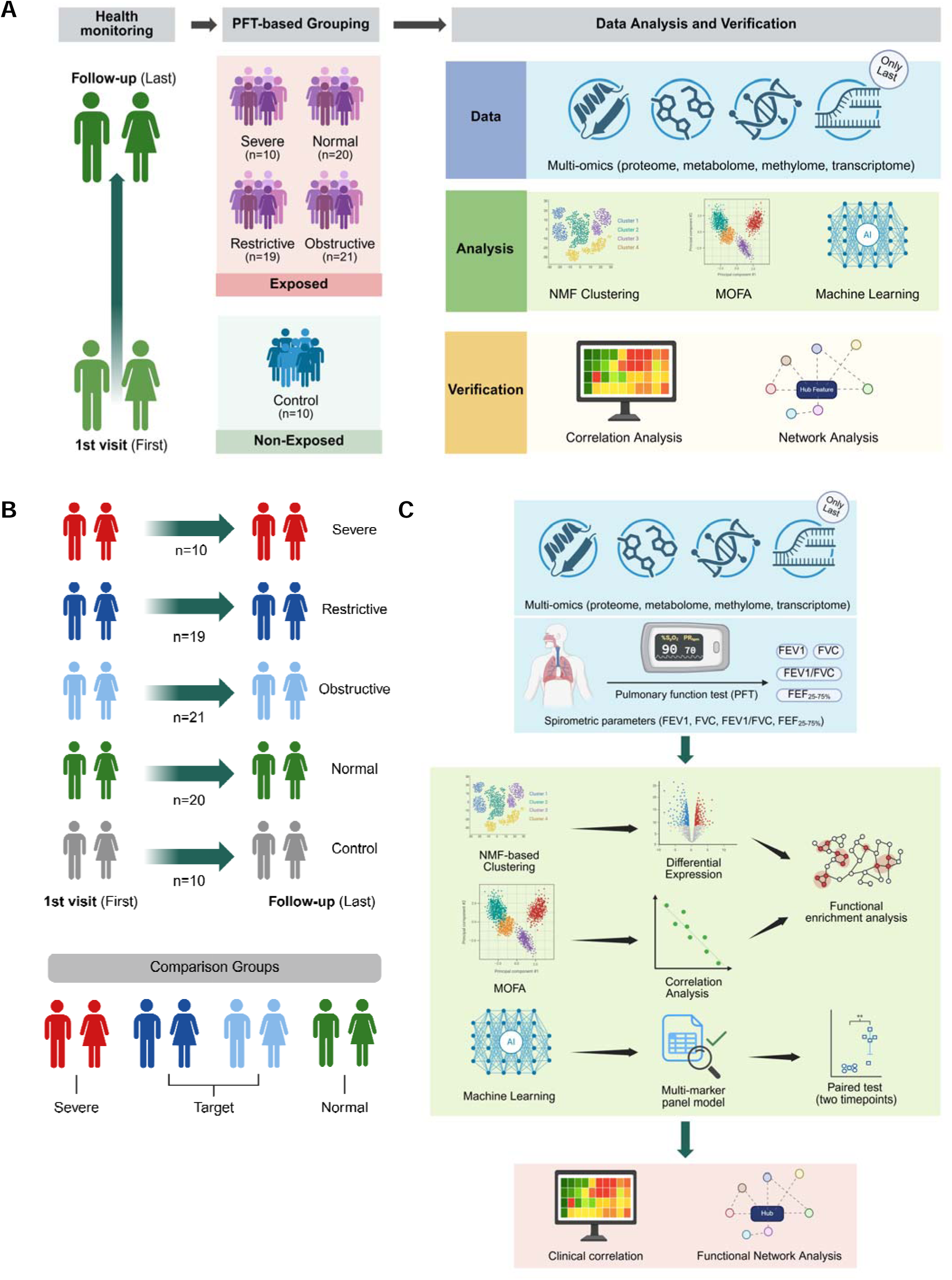
Overview of longitudinal assessment and analytical workflow. (A) The analytical framework comprises three main phases. First, all patients underwent health monitoring at Asan Medical Center and were stratified into four comparative groups and one control group: Severe (severe lung injury, n=10), Restrictive (restrictive lung disease, n=19), Obstructive (obstructive lung disease, n=21), Normal (normal lung function, n=20), and Control (control cohort, n=10). Second, comparative analysis was performed using multi-omics data and pulmonary function parameters through three key bioinformatic frameworks: NMF clustering, MOFA, and machine learning. Third, identified biomarker candidates were verified through correlation analysis to determine clinical associations and network analysis to infer network-level interactions. (B) The visualization displays paired longitudinal data from individuals within predefined clinical groups and designed comparison groups based on clinical parameters. (C) The analytical workflow illustrates the directional flow of this study. Multi-omics and spirometric parameters were integrated for comparative analysis and verification of biomarker candidates. Green arrows indicate the sequential order of each analytical phase, while black arrows represent the directional flow within individual data analysis steps.

**Table 1.** Primary characteristics of the study subjects.

| Characteristics | Unexposed-normal control (n=10) | Exposed-normal control (n=20) | Obstructive (n=21) | Restrictive (n=19) | Severe (n=10) | P-value |
| --- | --- | --- | --- | --- | --- | --- |
| 1st age (years) | 8.7±0.5 | 8.4±2.7 | 8.6±2.5 | 9.4±3.0 | 9.7±3.1 | 0.580* |
| Women, n (%) | 7(70.0) | 9 (45.0) | 10 (47.6) | 11 (57.9) | 3 (30.0) | 0.417** |
| PFT (1st) |  |  |  |  |  |  |
| Z-FVC | 0.0±1.4 | 0.8±1.2 | 0.0±1.4 | -2.1±0.9 | -2.8±0.6 | 0.000* |
| Z-FEV<sub>1</sub> | 0.2±0.9 | 1.0±0.9 | -0.3±1.1 | -1.5±0.8 | -2.9±0.5 | 0.000* |
| Z-FEV<sub>1</sub>/FVC | 0.3±1.0 | 0.5±0.9 | -0.6±1.0 | 0.7±0.9 | -0.8±1.4 | 0.000* |
| Z-FEF25-75% | 0.5±0.7 | 1.2±0.7 | -0.5±0.9 | 0.3±0.9 | -1.8±1.0 | 0.000* |
| PFT (4th) |  |  |  |  |  |  |
| Z-FVC | -0.5±1.1 | 0.5±0.6 | 0.7±1.3 | -1.6±0.7 | -2.6±1.3 | 0.000* |
| Z-FEV<sub>1</sub> | -0.2±1.3 | 0.9±0.6 | -0.4±1.1 | -1.1±0.7 | -3.0±1.0 | 0.000* |
| Z-FEV<sub>1</sub>/FVC | 0.3±1.0 | 0.8±1.3 | -1.6±0.8 | 0.4±1.0 | -1.5±1.3 | 0.000* |
| Z-FEF25-75% | 0.1±1.2 | 0.9±0.9 | -1.2±0.8 | 0.2±0.9 | -2.3±1.1 | 0.000* |
| Exposure concentration (ug/m<sup>3</sup>) | 0 | 117.5±84.0 | 86.4±80.2 | 148.6±146.5 | 179.4±150.0 | 0.002* |
| Accumulated usage time(hr) | 0 | 4644.8±4969.7 | 5887.6±9454.5 | 4242.4±6278.4 | 5306.0±3509.1 | 0.204* |
| Accumulated exposure levels | 0 | 486408.9±485755.5 | 342304.0±377798.3 | 435102.0±433268.8 | 949457.1±1085445.4 | 0.004* |

PFT, pulmonary function test; FVC, forced vital capacity; FEV<sub>1</sub>, forced expiratory volume in 1 second; FEF25-75%, forced expiratory flow at 25%-75% of FVC; \*ANOVA p-value, \*\*Chi-square p-value

### K-means Clustering

We initially evaluated whether conventional clustering techniques could effectively discriminate patients in a manner consistent with established clinical classifications. To evaluate the capacity of conventional clustering approaches to discriminate between clinical groups based on pulmonary function parameters, K-means clustering was implemented as the primary analytical strategy. The optimal number of clusters was determined through elbow method analysis, with subsequent visualization via Principal Component Analysis (PCA) to assess the method’s discriminatory capability across predefined clinical groups. This analytical framework was applied independently to both the initial (First) and final (Last) timepoint datasets. PCA projections demonstrated limited concordance between the computationally derived clusters and the established clinical group classifications (Figures E1A and E1B). Optimization procedures incorporating both elbow method and silhouette analysis converged on *k* = 4 as the optimal cluster number for both datasets (Figure E1C). To further characterize the clustering structure, hierarchical clustering using Ward linkage was performed on the K-means clustering results, with the resulting dendrogram (Figure E1D) corroborating the insufficient separation of clinical groups through this conventional approach. These findings reveal that K-means clustering, despite its widespread application in biomedical data analysis, exhibits limited efficacy in capturing the inherent heterogeneity among clinical groups when applied to our integrated multi-omics and clinical dataset.

### Non-negative Matrix Factorization (NMF)-based Clustering

In response to the limitations observed with conventional clustering approaches, we implemented Non-negative Matrix Factorization (NMF) as an alternative analytical strategy. NMF’s inherent non-negativity constraints confer distinct advantages in feature extraction and biological interpretability, particularly suited for omics data analysis where expression values are naturally non-negative (16). The NMF-based clustering was performed on the integrated omics dataset using the NMF package in R (29). Optimal factorization rank determinization was achieved through systematic evaluation of multiple metrics. Cophenetic correlation coefficients and dispersion values were computed across a range of factorization ranks, with the optimal rank identified at the inflection point where the rate of cophenetic score decrease exhibited marked acceleration relative to dispersion changes (Figure 2A). Consensus maps were subsequently generated to assess the stability and robustness of cluster assignments. Based on these comprehensive evaluations, rank = 4 was selected for the initial timepoint dataset and rank = 5 for the final timepoint dataset. To evaluate the clinical relevance of the NMF-derived clusters, we examined the distribution of Pulmonary Function Test (PFT) parameters across the identified subgroups (Figure 2B). The PFT values, provided as clinical metadata, were visualized using boxplots to characterize the functional respiratory profiles associated with each cluster. For subsequent comparative analyses, clusters were stratified based on z-score normalized PFT measurements. Clinical risk stratification was performed according to established pulmonary function criteria (24), with a z-score threshold of < -1.64 (corresponding to the 5^th^ percentile) designating the high-risk group. This classification yielded the following comparison groups: for the initial dataset, Cluster 3 (high-risk) vs Cluster 1 (control); for the final dataset, Cluster 3 (high-risk) vs Cluster 1 (control). The NMF-based approach demonstrated superior capability in delineating clinically meaningful subgroups, revealing distinct molecular patterns associated with differential pulmonary function outcomes.

**Figure 2.**
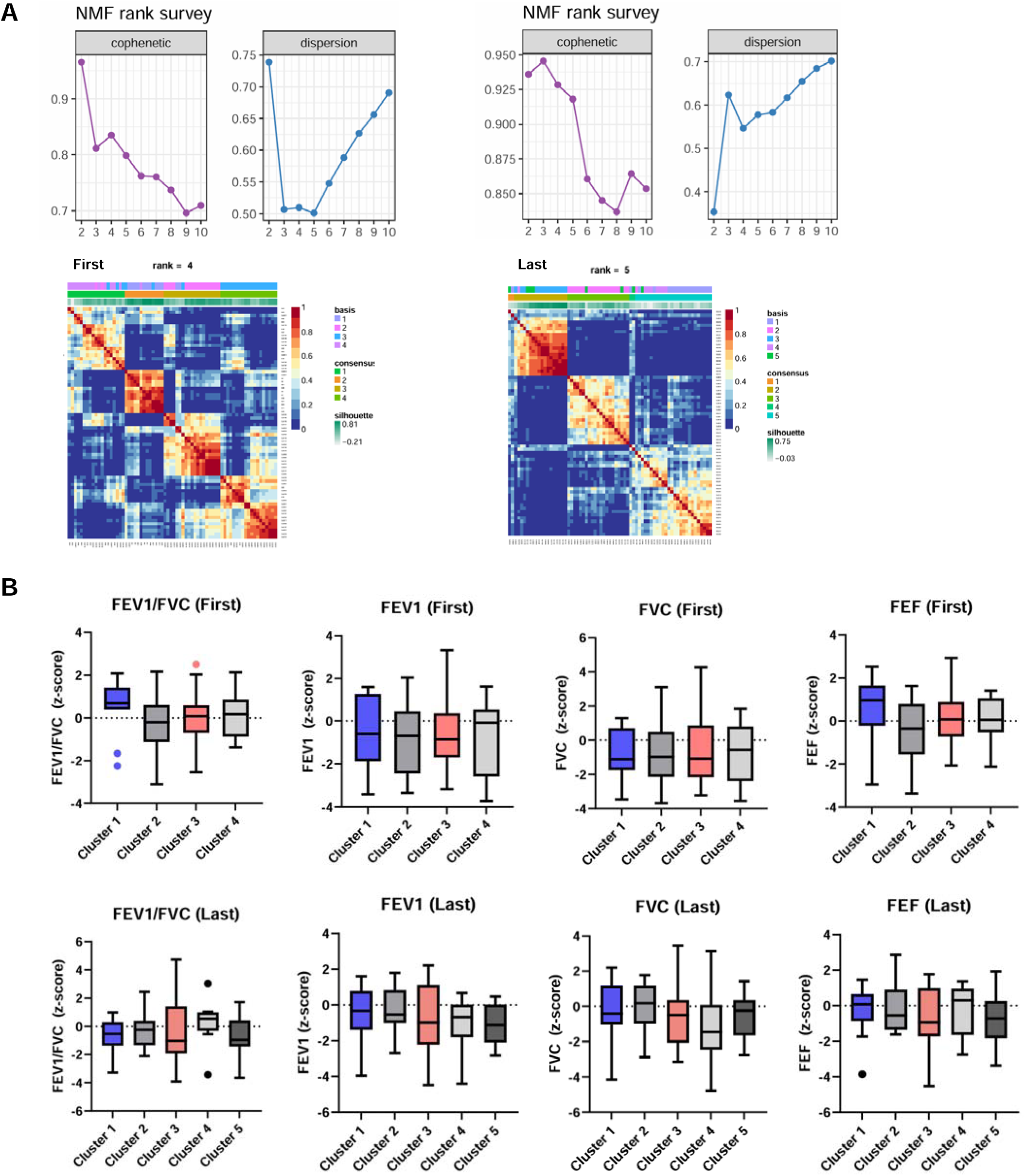
Non-negative Matrix Factorization clustering effectively discriminates patient subgroups based on pulmonary function profiles. (A) Determination of optimal factorization rank for NMF-based clustering analysis. Top panels display cophenetic correlation coefficients and dispersion values across candidate ranks for initial and final timepoint datasets. Optimal ranks were identified at the inflection points where cophenetic scores exhibited marked decline (rank = 4 for initial dataset; rank = 5 for final dataset). Bottom panels provide comprehensive cluster quality assessments through basis matrix visualization, consensus maps, and silhouette coefficient distributions, confirming robust cluster assignment and stability at the selected ranks. (B) Distribution of spirometry-derived Pulmonary Function Test (PFT) parameters across NMF-identified clusters. Box plots demonstrate the stratification of FEV₁, FVC, and FEV₁/FVC ratio values within each cluster for both timepoints. Clusters characterized by impaired pulmonary function (z-score < -1.64, meeting clinical criteria for lung function abnormality) are highlighted in salmon red, whereas clusters exhibiting preserved lung function (control groups) are depicted in royal blue. The distinct separation of PFT distributions between high-risk and control clusters validates the clinical relevance of the NMF-based stratification approach.

### Molecular Characterization of NMF-Derived Clusters

To investigate the molecular signatures underlying the NMF-stratified patient subgroups, we performed comprehensive differential expression analyses across multiple omics layers. Initial transcriptomics analysis using DESeq2 revealed differential gene expression between clusters, with insufficient differentially expressed genes (DEGs) meeting our stringent criteria (fold change > 1.5, adjusted p-value < 0.05) to support robust Gene Set Enrichment Analysis (GSEA). Given the limited transcriptomics signal, we redirected our analytical focus to the proteomic level, employing the DEP package (31) for differential protein expression analysis. This approach successfully identified numerous differentially expressed proteins (DEPs) meeting our statistical thresholds (fold change > 1.5, adjusted p-value < 0.05) between high-risk group and control group (Figure 3A). The six most significantly altered proteins, ranked by combined log_2_ fold change and -log_10_ p-value, demonstrated clear separation between cluster phenotypes when their log_2_-centered intensities were visualized as bar plots (Figure 3B). Functional enrichment analysis of the identified DEPs was performed using g:Profiler, interrogating Gene Ontology (GO), Kyoto Encyclopedia of Genes and Genomes (KEGG), and Reactome databases. The directionality of pathway enrichment was assessed relative to the high-risk cluster designation. For the final dataset, pathway analysis revealed significant enrichment in several biological processes relevant to pulmonary pathophysiology (Figure 3C). Parallel analysis of the initial dataset identified proteins specifically enriched in lung injury-associated pathways, with their differential expression patterns visualized across clusters (Figure 3D). Notably, integrin-mediated signaling pathways exhibited opposite directions of regulation at the two timepoints. Key protein candidates emerged from this analysis, including PF4, CD36, PPBP, FERMT3, and RTN4 (Uniprot: Q9NQC3, isoform 3) in the final dataset, which participate in platelet activation, integrin signaling, and immune response pathways. Similarly, LIMS1 (Uniprot: P48059, isoform 2) and MYH3 in the initial dataset showed enrichment in integrin-related processes, with all candidates appearing in at least two independent pathway analyses (Table 2). These findings demonstrate that NMF-based stratification, when integrated with multi-omics profiling, can effectively identify molecular biomarkers associated with differential pulmonary function outcomes. However, we acknowledge inherent limitations in clustering methodologies, including the potential for artifactual cluster identification in the absence of true biological substructures (36) and the influence of investigator bias in cluster interpretation (37).

**Figure 3.**
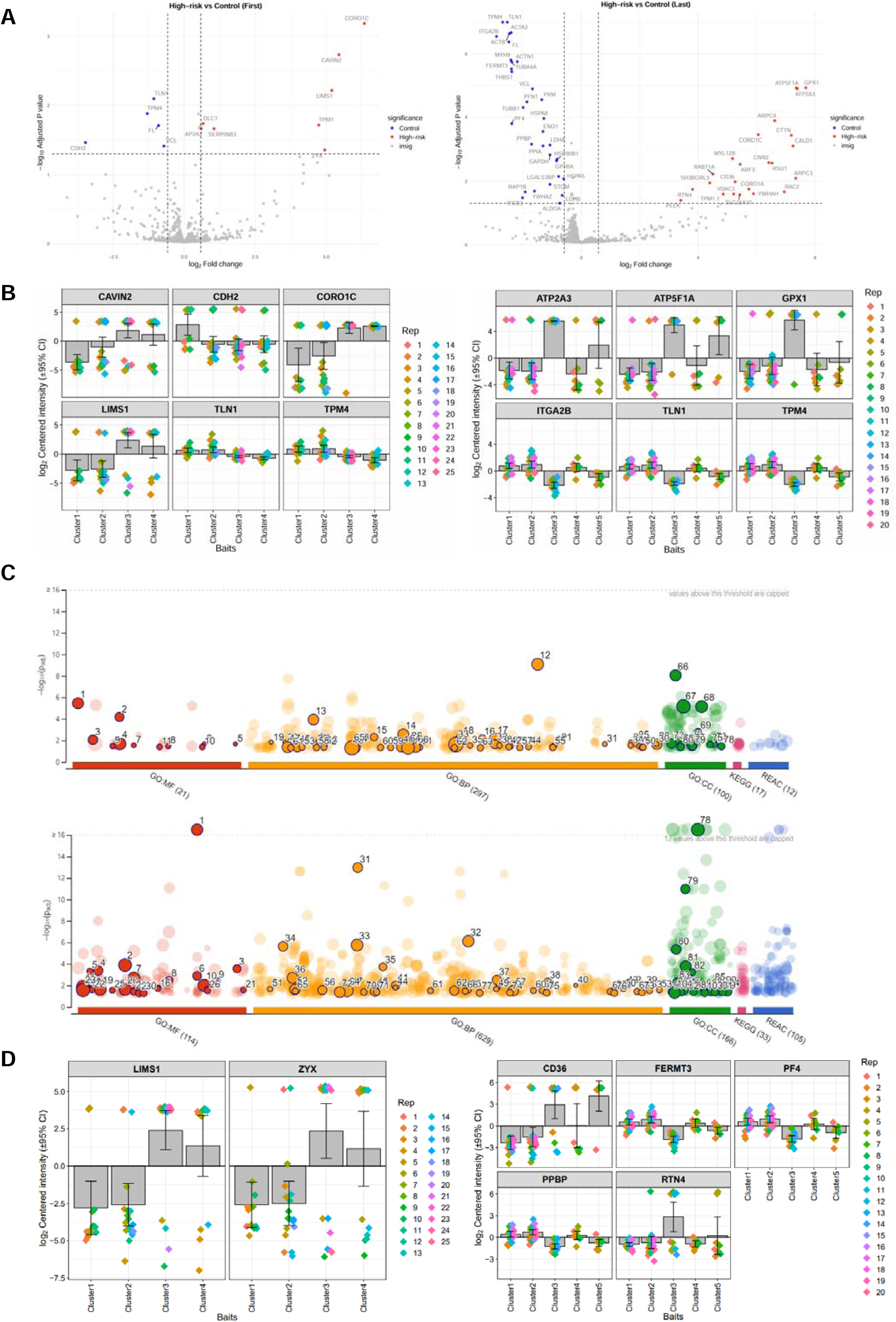
Proteomic profiling reveals differential molecular signatures between NMF-stratified patient clusters. (A) Volcano plots depicting differential protein expression analysis between high-risk and control clusters for initial and final timepoint datasets. Protein meeting significance criteria (|log_2_ fold change| > 1.5, adjusted p-value < 0.05) are highlighted in red (upregulated) and blue (downregulated). Gray dots represent proteins not meeting statistical thresholds. (B) Box plots visualization of log_2_-transformed protein intensities for the six most significantly altered proteins from each dataset at initial (left) and final (right) timepoints. Rows represent individual proteins ranked by statistical significance, and columns represent individual samples organized by cluster assignment. (C) Functional enrichment analysis of differential expressed proteins in the final dataset using g:Profiler. Bar plots display significantly enriched pathways (Benjamini-Hochberg FDR < 0.05) from Gene Ontology, KEGG, and Reactome databases, separated by upregulated (top) and downregulated (bottom) protein sets. Bar length indicates -log_10_(adjusted p-value), with pathway names ordered by statistical significance. (D) Box plots illustrating the distribution of log_2_-centered protein intensities for candidate biomarkers associated with lung injury pathways at initial (left) and final (right) timepoints. Proteins shown demonstrated consistent differential expression between clusters and enrichment in relevant biological processes.

**Table 2.** Differentially Expressed Proteins Enriched in Pulmonary Pathophysiology-Realated Pathways Following NMF-Based Stratification.

| Index | Pathway | Term_id | Timepoint | Regulation | Adjusted p-value | Proteins |
| --- | --- | --- | --- | --- | --- | --- |
| 1 | Integrin-mediated signaling pathway | GO:0007229 | first | up | 0.008 | LIMS1,MYH3 |
| 2 | Positive regulation of execution phase of apoptosis | GO:1900119 | first | up | 0.018 | DLC1 |
| 3 | Cellular response to transforming growth factor beta stimulus | GO:0071560 | first | up | 0.019 | LIMS1,MYH3 |
| 4 | Lipoteichoic acid immune receptor activity | GO:0070892 | last | up | 0.000 | CD36 |
| 5 | Positive regulation of macrophage cytokine production | GO:0060907 | last | up | 0.014 | CD36,RTN4 |
| 6 | Negative regulation of apoptotic signaling pathway | GO:2001234 | last | up | 0.021 | CTTN,GPX1,SLC25A31 |
| 7 | Positive regulation of pattern recognition receptor signaling pathway | GO:0062208 | last | up | 0.022 | CD36,RTN4 |
| 8 | Regulation of cytoplasmic pattern recognition receptor signaling pathway | GO:0039531 | last | up | 0.031 | CD36,RTN4 |
| 9 | CXCR chemokine receptor binding | GO:0045236 | last | down | 0.000 | PF4,PPBP |
| 10 | Platelet activation | GO:0030168 | last | down | 0.010 | ACTB,FERMT3,FLNA,GP1BA,ITGB3,MYH9,PF4,PPIA,T<br>LN1,TUBB1,VCL |
| 11 | Integrin activation | GO:0033622 | last | down | 0.015 | FERMT3,RAP1B,TLN1 |
| 12 | Antimicrobial humoral immune response mediated by antimicrobial peptide | GO:0061844 | last | down | 0.015 | GAPDH,PF4,PPBP |
| 13 | Cellular response to transforming growth factor beta stimulus | GO:0071560 | last | down | 0.026 | ACTA2,HSPAS,THBS1 |
| 14 | Platelet alpha granule | GO:0031091 | last | down | 0.034 | ACTN1,ALDOA,FERMT3,ITGA2B,ITGB3,PF4,PPBP,TH<br>BS1 |

### Multi-Omics Factor Analysis Reveals Molecular Signatures Associated with Pulmonary Function

To overcome the inherent limitations of clustering-based approach, we implemented Multi-Omics Factor Analysis (MOFA), an unsupervised learning framework that enables simultaneous integration of multiple omics layers with clinical metadata (18). Unlike NMF, MOFA employs a Bayesian probabilistic model as dimensionality reduction strategy to identify latent factors that capture sources of variation across heterogeneous data types (17). The MOFA model was applied independently to both initial and final timepoint datasets (Figure 4A). The analysis yielded 9 factors for the initial dataset and 11 factors for the final dataset, with each factor explaining a distinct proportion of variance across one or more data views. To assess the relative contribution of each omics layer, we quantified the proportion of variance explained by metabolomic, methylomic, and proteomic data across all identified factors and clinical groups, with transcriptomic data included exclusively in the final timepoint dataset (Figure 4B). Association analysis between MOFA factors and clinical parameters revealed significant correlations with pulmonary function metrics and demographic variables. Specifically, two factors (factors 2 and 6) demonstrated strong associations with PFT parameters at both timepoints, while age showed broader associations across four distinct factors (factors 3, 6, 8, and 10). Of particular interest, factor 6 exhibited significant correlations with FEV_1_/FVC ratio and forced expiratory flow (FEF) specifically within the Target group, while maintaining its association with age across all clinical groups (Figure 4C). Molecular characterization of Factor 6 revealed Collagen alpha-1(XVIII) chain as the protein with the highest loading weight among the top 10 contributors (Figure 4D). Functional enrichment analysis was performed separately on proteins with positive and negative loadings. Positively weighted proteins showed significant enrichment in integrin cell surface interactions (highest -log_10_ p-value), alongside signaling cascades and immune response pathways. In contrast, negatively weighted proteins were predominantly enriched in positive regulation of lipid metabolism and related metabolic processes (Figure 4E). Remarkably, integrin-mediated pathways emerged as consistently upregulated across both MOFA and NMF analyses, reinforcing their potential role in pulmonary dysfunction. Through systematic evaluation of protein loadings and pathway enrichment frequencies, we identified key biomarker candidates: A2M and ACTN1 (positive loadings), and ABCA1, ADIPOQ, AGT, APOA1, and APOA2 (negative loadings), all appearing in ≥ 7 enriched pathways among the top 10 weighted proteins. ACTB, despite high frequency in both positive and negative loadings, was excluded due to its ubiquitous expression and lack of specificity (Table 3).

**Figure 4.**
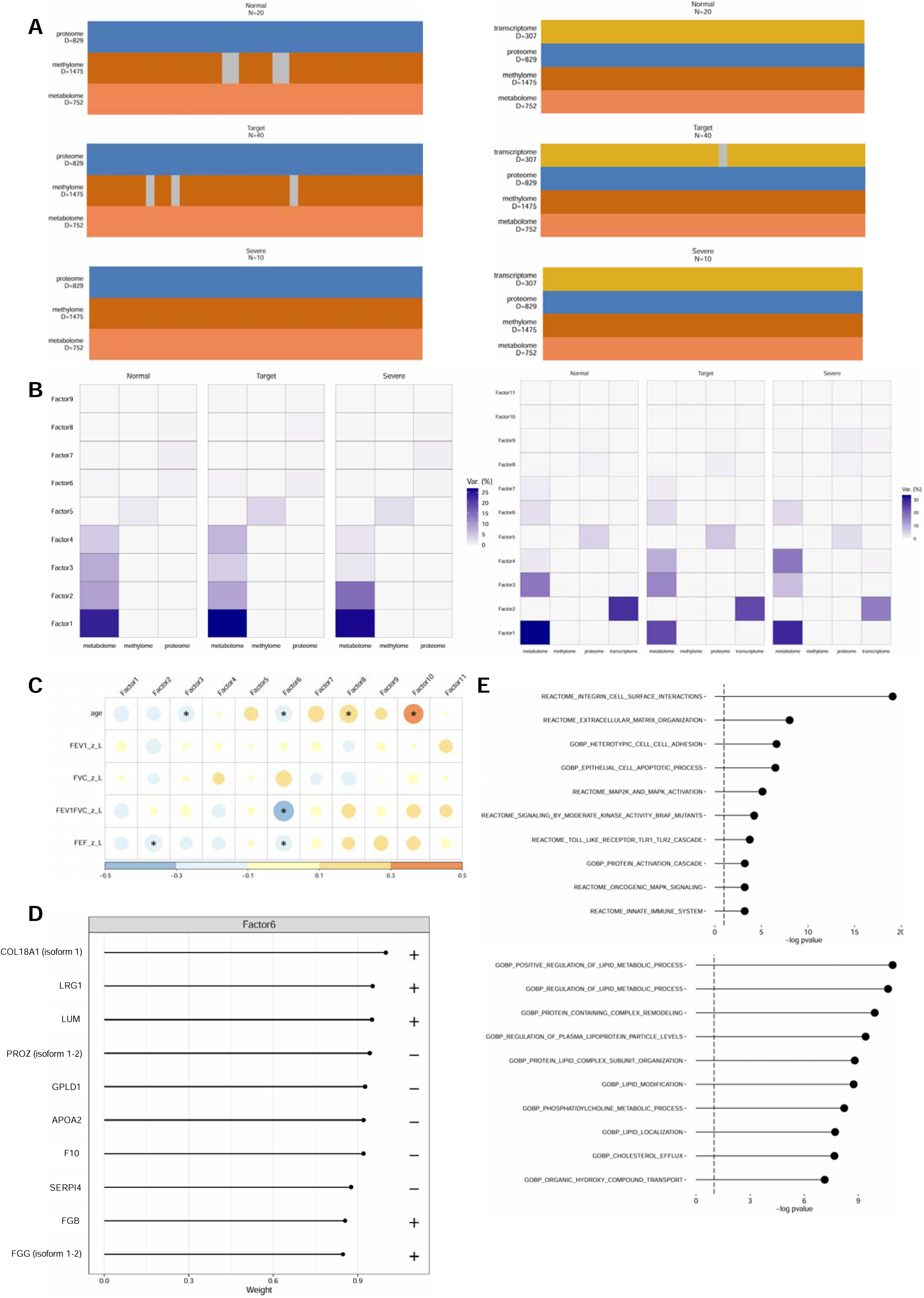
Multi-Omics Factor Analysis reveals target group-specific molecular signatures associated with pulmonary function decline. (A) Overview of Multi-Omics Factor Analysis (MOFA) implementation showing data structure and clinical group stratification for initial (left) and final (right) timepoint datasets. Groups are defined as: Normal (normal lung function), Target (target population with mild impairment), and Severe (severe lung injury). (B) Variance decomposition analysis displaying the relative contribution of metabolomic, methylomic, and proteomic data layers to each latent factor, with transcriptomic data included exclusively in the final timepoint dataset. MOFA identified 9 factors in the initial dataset and 11 factors in the final dataset, each explaining variance of all views. (C) Heatmap depicting Pearson correlation coefficients between MOFA-derived factors and clinical parameters in the final dataset. Significant associations (p < 0.05) are marked with asterisks. Factor 6 shows specific correlations with FEV1/FVC and FEF measurements in the Target. (D) Protein loading weights for Factor 6, displaying the top 10 contributors ranked by absolute weight values. Collagen alpha-1(XVIII) chain exhibits the highest loading, indicating its strong association with the pulmonary function-related variance captured by this factor. (E) Functional enrichment analysis of Factor 6-associated proteins, separated by loading direction. Upper panel shows pathways enriched among positively weighted proteins, with integrin cell surface interactions displaying the highest significance. Lower panel depicts pathways enriched among negatively weighted proteins, dominated by lipid metabolism processes. Bar length represents -log_10_(adjusted p-value), with pathway selection based on Benjamini-Hochberg FDR < 0.05.

**Table 3.** Factor 6-Associated Proteins with High Pathway Enrichment Frequencies Identified through MOFA-Driven Functional Analysis.

| Index | Pathway | Timepoint | Regulation | P-value | Size | Genes |
| --- | --- | --- | --- | --- | --- | --- |
| 1 | REACTOME_INTEGRIN_CELL_SURFACE_INTERACTIONS | last | positive | 8.30E-20 | 25 | TTR, HSPG2, APOB, AHSG, COL6A3, ALDOA, COL18A1, ITGA2B, VWF, C4B_2, ITGA6, THBS1, FGG, FN1, ICAM1, COMP, COL10A1, ITGB3, VCAM1, COL6A1 |
| 2 | REACTOME_EXTRACELLULAR_MATRIX_ORGANIZATION | last | positive | 9.50E-09 | 48 | TTR, HSPG2, APOB, AHSG, COL6A3, ALDOA, A2M, COL18A1, EFEMP1, TNXB, ITGA2B, VWF, C4B_2, ITGA6, MMP2, PLG, NCAM1, THBS1, FGG, ACTN1 |
| 3 | GOBP_HETEROTYPIC_CELL_CELL_ADHESION | last | positive | 2.40E-07 | 11 | APOA1, FGB, FGG, CD44, FGA, ITGB3, VCAM1, ADIPOQ, NRCAM, ITGB1, SIRPA |
| 4 | GOBP_EPITHELIAL_CELL_APOPTOTIC_PROCESS | last | positive | 3.20E-07 | 10 | GSN, THBS1, FGB, FGG, FGA, ICAM1, CDH5, KRT18, PKHD1, BAD |
| 5 | REACTOME_MAP2K_AND_MAPK_ACTIVATION | last | positive | 7.60E-06 | 11 | TLN1, VCL, ITGA2B, VWF, ACTB, FGB, FGG, FGA, FN1, ITGB3, RAP1B |
| 6 | REACTOME_SIGNALING_BY_MODERATE_KINASE_ACTIVITY_BRAF_MUTANTS | last | positive | 5.90E-05 | 12 | TLN1, VCL, ITGA2B, VWF, ACTB, FGB, FGG, FGA, FN1, ITGB3, CALM1, RAP1B |
| 7 | REACTOME_TOLL_LIKE_RECEPTOR_TLR1_TLR2_CASCADE | last | positive | 1.70E-04 | 10 | CD14, FGB, FGG, S100A8, FGA, S100A9, SAA1, UBB, MAPKAPK3, CD36 |
| 8 | GOBP_PROTEIN_ACTIVATION_CASCADE | last | positive | 5.90E-04 | 14 | SERPINC1, F12, FGB, FGG, GP9, FGA, FN1, APOH, F13A1, F13B, KLKB1, GP5, GP1BA, GP1BB |
| 9 | REACTOME_ONCOGENIC_MAPK_SIGNALING | last | positive | 6.10E-04 | 14 | TLN1, VCL, ITGA2B, VWF, ACTB, FGB, FGG, FGA, FN1, ITGB3, CALM1, RAP1B, AKAP9, CLCN6 |
| 10 | REACTOME_INNATE_IMMUNE_SYSTEM | last | positive | 6.10E-04 | 190 | C4A, TTR, APOB, AHSG, IGHE, QSOX1, ALDOA, VCL, C7, C3, CFI, PIGR, MYH9, HSP90AA1, IGLC2, CAT, KRT1, C8G, IGHV3-23, IGHV3-7 |
| 11 | GOBP_POSITIVE_REGULATION_OF_LIPID_METABOLIC_PROCESSES | last | negative | 1.20E-11 | 10 | AGT, APOE, APOA1, F2, GPLD1, APOA4, APOC1, APOA2, APOC2, ADIPOQ |
| 12 | GOBP_REGULATION_OF_LIPID_METABOLIC_PROCESS | last | negative | 2.20E-11 | 18 | APOB, AGT, APOE, C3, APOA1, F2, GPLD1, APOA4, APOD, APOC3, APOC1, SNCA, APOA2, APOC2, ADIPOQ, H6PD, RARRES2, PIBF1 |
| 13 | GOBP_PROTEIN_CONTAINING_COMPLEX_REMODELING | last | negative | 1.20E-10 | 13 | APOB, AGT, LCAT, APOE, APOA1, PLTP, APOA4, CETP, APOC3, APOC1, APOA2, APOC2, PON1 |
| 14 | GOBP_REGULATION_OF_PLASMA_LIPOPROTEIN_PARTICLE_LEVELS | last | negative | 3.90E-10 | 19 | APOB, AGT, LCAT, APOE, APOA1, PLTP, KHSRP, PCSK9, GPLD1, APOA4, CETP, APOC3, APOC1, APOA2, APOC2, PON1, ADIPOQ, ABCA1, CD36 |
| 15 | GOBP_PROTEIN_LIPID_COMPLEX_SUBUNIT_ORGANIZATION | last | negative | 1.60E-09 | 14 | APOB, AGT, LCAT, APOE, APOA1, PLTP, APOA4, CETP, APOC3, APOC1, APOA2, APOC2, PON1, ABCA1 |
| 16 | GOBP_LIPID_MODIFICATION | last | negative | 1.80E-09 | 10 | AGT, LCAT, APOE, APOA1, APOA4, APOD, APOC1, APOA2, ADIPOQ, SACM1L |
| 17 | GOBP_PHOSPHATIDYLCHOLINE_METABOLIC_PROCESS | last | negative | 6.00E-09 | 11 | LCAT, APOA1, GPLD1, APOA4, CETP, APOC1, APOA2, PON1, PNPLA8, CHKB, ENPP2 |
| 18 | GOBP_LIPID_LOCALIZATION | last | negative | 1.90E-08 | 40 | SLC4A1, APOB, AGT, LCAT, APOE, C3, APOA1, PLTP, THBS1, PCSK9, APOL1, CLU, APOA4, CETP, CRP, LBP, APOH, APOC4, CFHR4, APOD |
| 19 | GOBP_CHOLESTEROL_EFFLUX | last | negative | 2.10E-08 | 14 | SLC4A1, APOE, C3, APOA1, PLTP, APOL1, APOA4, CETP, CRP, APOH, APOC3, APOC1, ATP10A, APOF |
| 20 | GOBP_ORGANIC_HYDROXY_COMPOUND_TRANSPORT | last | negative | 7.30E-08 | 25 | SLC4A1, APOB, APOE, C3, APOA1, PLTP, ACTB, PCSK9, APOL1, PARK7, APOA4, CETP, CRP, LBP, APOH, APOC4, ITGB3, APOC3, APOC1, SNCA |

### Machine Learning-Based Multi-Biomarker Panel Development

To address the limitations of single biomarker approaches and leverage the complementary information from multi-omics profiling, we applied a machine learning framework for multi-biomarker panel identification (19, 20). Using RapidMiner Studio v9.10.010, we systematically evaluated 18 distinct machine learning algorithms on the initial timepoint dataset. Model performance was assessed through 3-fold cross-validation to ensure robust generalization estimates. Among the tested algorithms, 12 achieved classification accuracy exceeding 70%, with 6 models, including deep learning architectures, demonstrating accuracy above 80% (Figure 5A). Feature importance analysis the 12 high-performing models (accuracy > 70%) revealed consistent selection patterns. IGHV2-70 emerged as the most frequently selected biomarker (frequency = 11/12 models), followed by LysoPC (16:0) (frequency = 3, confidence level = 2) and hexadecyl ferulate (frequency = 2, confidence level = 3). These markers aligned with the features prioritized through forward selection in the primary model (Figure 5B). The deep learning model demonstrated robust discriminatory performance among the three clinical groups (Normal, Target, Severe), achieving 81.46% accuracy on the initial timepoint dataset (Figure 5C). Receiver operating characteristic (ROC) analysis revealed strong discriminatory power, with the area under the curve (AUC) for the Target reaching 0.779, indicating good sensitivity-specificity balance (Figure 5D). To validate the temporal stability of the identified biomarkers, we performed longitudinal analysis comparing initial and final timepoint measurements. Wilcoxon signed-rank tests demonstrated significant temporal correlations for IGHV2-70, LysoPC (16:0), and hexadecyl ferulate (p < 0.05), confirming their consistent association with disease progression. In contrast, CLCN6 (Uniprot: P51797, isoform 5) and SPTBN5 showed no significant temporal correlation, suggesting potential state-dependent variations (Figure 5E). Based on these comprehensive analyses, we propose a multi-biomarker panel consisting of IGHV2-70, LysoPC (16:0), and hexadecyl ferulate.

**Figure 5.**
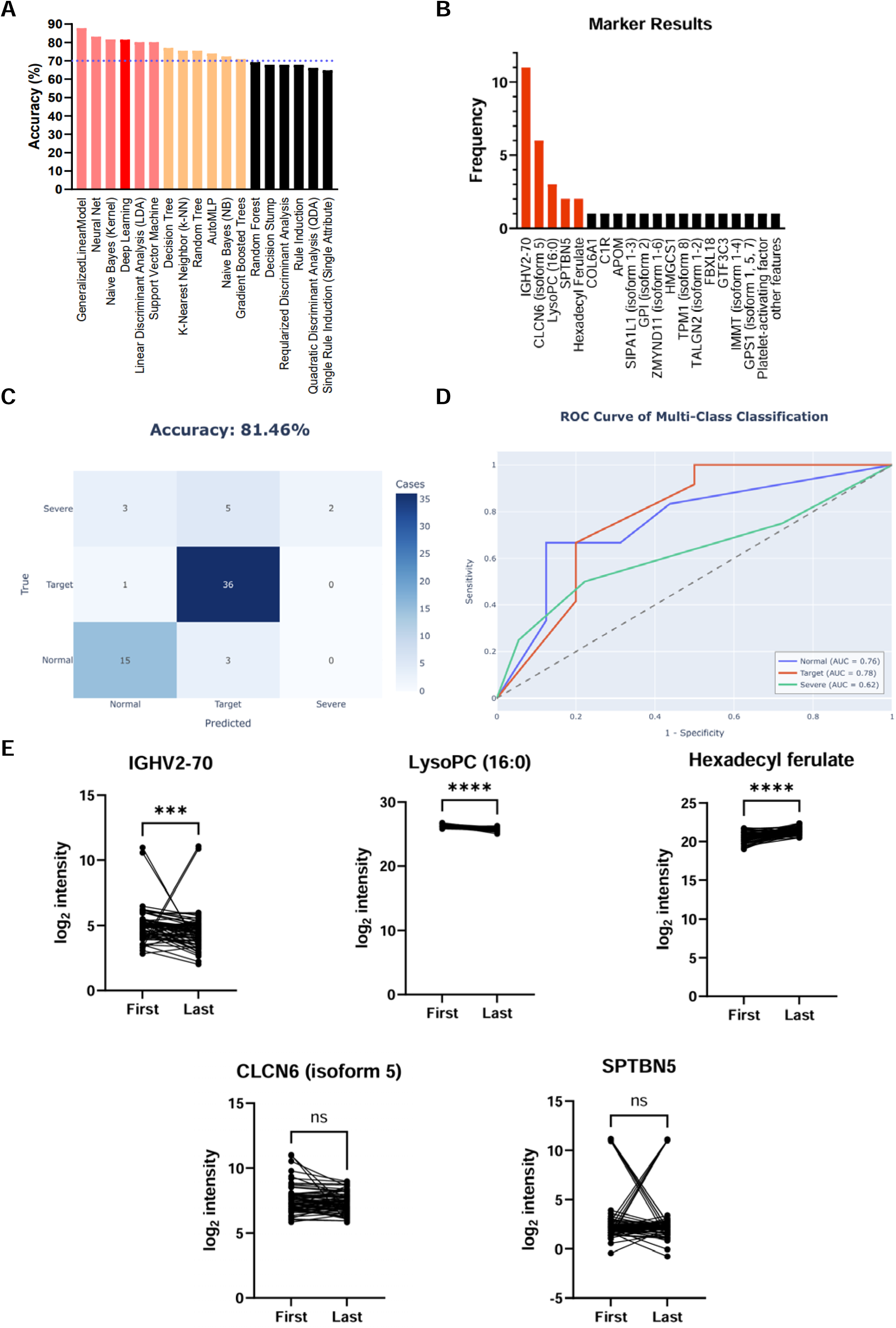
Machine learning-based identification of a multi-biomarker panel for clinical stratification. (A) Comparative performance evaluation of 18 machine learning algorithms applied to the initial timepoint dataset with 3-fold cross-validation. Model accuracies are color-coded: beige (70-80%) and salmon red (>80%). Deep Learning achieved the highest performance at 81.46% accuracy (highlighted in red). Additional high-performing models include Linear Discriminant Analysis (LDA) and Support Vector Machine (SVM). (B) Feature selection frequency analysis across all 18 models during cross-validation. Bar height indicates the number of models selecting each biomarker. Features selected by more than two models are highlighted in red, with IGHV2-70 showing universal selection (frequency = 11/12). (C) Confusion matrix demonstrating the Deep Learning model’s classification performance across three clinical groups. True labels (y-axis) versus predicted labels (x-axis) show the model’s ability to discriminate between Normal, Target, and Severe populations, achieving 81.46% overall accuracy. (D) Receiver Operating Characteristic (ROC) curves for multi-class classification using Deep Learning-derived probability scores. Area Under the Curve (AUC) values: Normal (0.755), Target (0.779), Severe (0.625), with macro-average AUC of 0.72, indicating good discriminatory power across all clinical groups. (E) Longitudinal analysis of selected biomarker intensities showing paired measurements between initial and final timepoints for individual patients. Lines connect measurements from the same patient. Statistical significance assessed using Wilcoxon signed-rank test for non-normally distributed data. Significant temporal stability observed for P01814, LysoPC (16:0), and hexadecyl ferulate (***p < 0.001, ****p < 0.0001), while P51797/P51797-5 and Q9NRC6 showed no significant correlation (ns: not significant).

### Clinical Correlation and Functional Network Analysis of Identified Biomarkers

To evaluate the clinical relevance of the multi-omics-derived biomarkers, we performed comprehensive correlation analyses with pulmonary function parameters and investigated their molecular interactions through network-based approaches. Correlation analysis revealed modest but consistent associations between specific biomarkers and lung function metrics. In the initial timepoint, Platelet glycoprotein 4 (CD36), identified through NMF analysis, demonstrated a weak negative correlation with FEV_1_ (r = -0.32, p < 0.05). At the final timepoint, MOFA-identified proteins, ABCA1 and apolipoprotein A2 (APOA2), exhibited slight negative correlations with FVC (r = -0.28 and -0.31, respectively, p < 0.05). These correlations were specifically observed within the target groups suggesting disease-specific associations (Figure 6A). To determine whether these biomarker-clinical associations were dependent on disease severity, we performed stratified correlation analyses across the three clinical groups. Strikingly, biomarkers that showed no significant correlation with lung function parameters in the normal group demonstrated progressively stronger associations as disease severity increased. Most notably, CD36 exhibited a disease stage-dependent correlation pattern: while displaying only weak association with FEV_1_ in the target groups (r = -0.32, p < 0.05), it showed a robust negative correlation with FEV_1_/FVC ratio in the severe group (r = -0.67, p < 0.05). Furthermore, no significant correlation was identified in normal group (Figures 6A and E2). This progressive enhancement of biomarker-clinical correlations suggests that the identified molecular signatures become increasingly coupled with pulmonary dysfunction as pathological changes accumulate.

**Figure 6.**
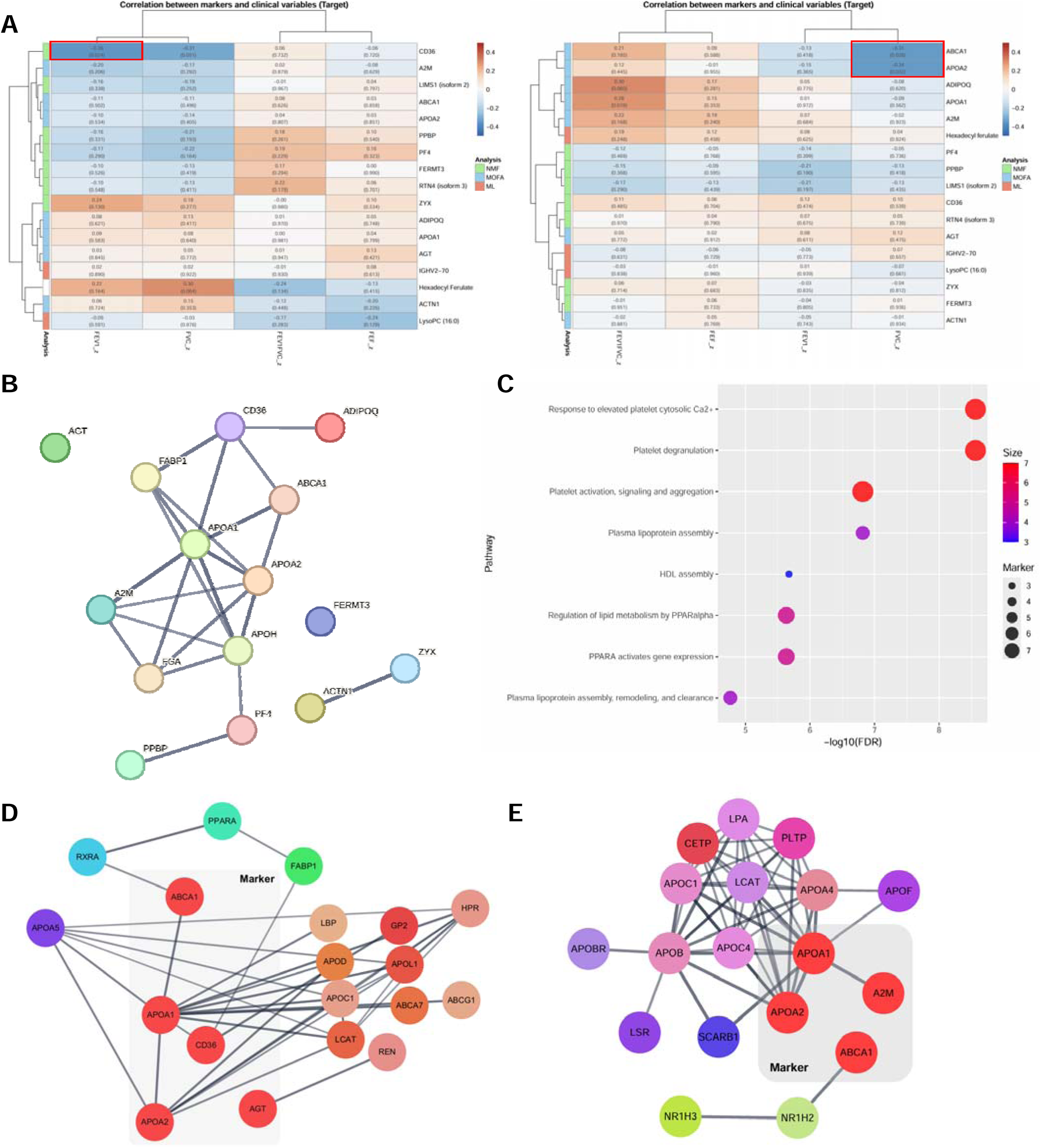
Clinical correlation and functional network analysis reveal biomarker convergence on pulmonary pathophysiology-related pathways. (A) Correlation heatmaps displaying associations between identified biomarkers and pulmonary function parameters in the target groups at initial (left) and final (right) timepoints. Statistically significant correlations (p < 0.05) are indicated by red squares. At the initial timepoint, CD36 shows negative correlation with FEV₁ (r = -0.32), while at the final timepoint, ABCA1 and APOA2 demonstrate negative correlations with FVC (r = -0.28 and -0.31, respectively). (B) Protein-protein interaction network constructed using STRING database (v11.5) with high confidence threshold (0.7). Network includes 12 of 17 biomarkers with maximum 3 interactors per node. Node size represents degree of connectivity, with edge thickness indicating interaction confidence. FERMT3 appears as an isolated node without direct interactions. (C) Pathway enrichment analysis using ReactomeFIViz displaying the top 8 pathways ranked by false discovery rate (FDR < 0.05). Bubble size represents the number of biomarkers involved in each pathway, while color intensity indicates statistical significance (-log_10_ FDR). Three major pathway clusters emerge: platelet activation, plasma lipoprotein metabolism, and lipid metabolic processes. (D) Functional interaction network for "Regulation of lipid metabolism by PPARalpha" pathway. Biomarkers (highlighted in coral red with grey borders) are shown within the context of their functional partners. Network expansion performed one step from seed proteins with confidence score 0.7. PPARA emerges as a central regulatory hub connecting ABCA1, APOA1, APOA2, and CD36. AGT shows isolated interaction with REN. (E) Functional interaction network for "Plasma lipoprotein assembly, remodeling, and clearance" pathway. Markov clustering (MCL) was applied to identify functionally coherent modules. Two clusters containing biomarkers (coral red nodes with grey borders) are displayed, revealing tight integration between lipoprotein metabolism (APOA1, APOA2, ABCA1) and inflammatory response (A2M) components.

To contextualize these biomarkers within broader molecular networks, we constructed protein-protein interaction networks using STRING database with high confidence threshold (0.7). The resulting network revealed extensive interconnectivity among most biomarkers with the notable exception of fermitin family homolog 3 (FERMT3), which appeared as an isolated node without direct interactions (Figure 6B). Functional enrichment analysis using ReactomeFIViz, a Cytoscape application for pathway visualization and analysis (38, 39), identified three major biological systems encompassing our biomarker panel: platelet activation and aggregation, plasma lipoprotein metabolism (particularly HDL-mediated processes), and lipid metabolic pathways (Figure 6C). Detailed network topology analysis revealed distinct functional modules. Within the lipid metabolism network, ABCA1, APOA1, APOA2, CD36, and AGT formed an interconnected cluster with PPARA emerging as a central regulatory hub. Notably, AGT demonstrated a unique interaction pattern, connecting exclusively with REN without direct associations to other network components (Figure 6D). The plasma lipoprotein assembly, remodeling, and clearance network incorporated APOA1, APOA2, A2M, and ABCA1. To enhance network interpretability, we applied Markov clustering (MCL) algorithm to identify functionally coherent submodules. Two distinct clusters containing our biomarkers were selected for detailed visualization, revealing tight functional coupling between lipoprotein metabolism and inflammatory response pathway (Figure 6E). These network analyses demonstrate that our biomarker panel converges on biologically coherent pathways previously implicated in pulmonary pathophysiology, particularly those governing lipid homeostasis, vascular integrity, and inflammatory responses. The observed clinical correlations, combined with the functional interconnectivity, support the biological relevance of these markers in lung injury progression among humidifier disinfectant lung injury.

## Discussion

This study employed an integrative multi-omics approach to identify molecular biomarkers associated with long-term pulmonary dysfunction in adolescents exposed to humidifier disinfectants. Through systematic application of multiple analytical frameworks, we uncovered distinct molecular signatures that stratify patients based on lung function outcomes and provide mechanistic insights into disease progression. Our initial clustering attempts using K-means methodology revealed fundamental limitations in capturing the biological heterogeneity of our patient cohort. The inability of conventional clustering to discriminate between clinical groups, as evidenced by overlapping PCA projections and poorly resolved dendrograms, necessitated more sophisticated approaches. This finding underscores a critical challenge in multi-omics analysis: the need for methods capable of handling non-linear relationships and complex interaction patterns inherent in biological systems.

The implementation of Non-negative Matrix Factorization (NMF) substantially improved patient stratification, enabling identification of clinically meaningful subgroups based on pulmonary function parameters. Proteomic profiling of NMF-derived clusters revealed compelling molecular signatures. In the initial timepoint, upregulated proteins converged on integrin-mediated signaling and apoptotic pathways. The prominence of integrin signaling aligns with its established role in TGF-β1 activation and pulmonary fibrosis pathogenesis (40). Notably, this finding corroborates our previous transcriptomic analysis in humidifier disinfectant-exposed children (14), suggesting persistent dysregulation of this pathway across multiple molecular layers.

The temporal evolution of molecular signatures revealed intriguing patterns. While integrin pathways dominated early responses, the final timepoint showed downregulation in immune response pathways, particularly platelet activation and chemokine signaling. This temporal shift may reflect the transition of immune modulation from platelet in chronic inflammatory states. Previous studies demonstrating platelets modulate immune responses through crucial interactions with integrins, cytokines, and chemokines (41). Remarkably, platelets can both activate immune cells to elicit immune responses and exert anti-inflammatory effects via cytokine secretion, particularly under inflammatory conditions (42). Proteomic analysis of patients with COPD revealed downregulation of platelet-mediated pathway proteins, including PF4, suggesting a potential impairment in recovery processes (43). Our analysis indicates temporal alterations in pulmonary immunity responses.

Multi-Omics Factor Analysis (MOFA) provided complementary insights by identifying latent factors capturing variance across multiple data modalities. Factor 6, specifically associated with FEV₁/FVC and FEF in the target group, revealed a striking dichotomy: positive loadings enriched in immune pathways (particularly integrin interactions), while negative loadings converged on lipid metabolism. This bipolar signature aligns with emerging evidence linking lipid metabolism dysregulation to idiopathic pulmonary fibrosis (44, 45), suggesting shared pathophysiological mechanisms between humidifier disinfectant exposure and other fibrotic lung diseases. The clinical relevance of our findings is underscored by the correlation patterns observed between biomarkers and pulmonary function parameters. CD36’s negative correlation with FEV₁, which intensified with disease severity, provides a potential monitoring tool for disease progression. Similarly, the negative correlations between ABCA1/APOA2 and FVC in the final dataset align with established associations between FVC decline and mortality in pulmonary fibrosis (46). The disease stage-dependent strengthening of these correlations suggests that molecular perturbations become increasingly coupled with functional decline as pathology advances.

Our machine learning framework successfully identified a multi-biomarker panel comprising IGHV-70, LysoPC (16:0), and hexadecyl ferulate, achieving 81.46% classification accuracy with an AUC of 0.779 for the target group. This performance level indicates clinically meaningful diagnostic potential (47). LysoPC (16:0), a key component of this panel, has established roles in airway inflammation (48) and metabolic crosstalk between adipose tissue and skeletal muscle (49). Given the documented muscle dysfunction in chronic respiratory diseases (50), this metabolite may represent a systemic marker of disease progression. The inclusion of hexadecyl ferulate, a poorly characterized metabolite, highlights the potential for discovering novel pathophysiological mechanisms through unbiased omics approaches.

Network analysis revealed that our biomarkers converge on three major biological systems: platelet activation, lipoprotein metabolism, and lipid homeostasis. The central role of PPARA in connecting multiple biomarkers suggests potential therapeutic targets. The tight integration between lipoprotein metabolism and inflammatory pathways, as revealed by functional interaction networks, provides a molecular framework for understanding the systemic nature of humidifier disinfectant-induced lung injury.

Several limitations warrant consideration. The temporal division of our dataset reduced statistical power for individual analyses, potentially limiting biomarker discovery. The minimal overlap between biomarkers identified through different analytical approaches reflects the complementary nature of these methods but also highlights the challenge of identifying universal markers. Additionally, the lack of an independent validation cohort necessitates cautious interpretation of our findings.

Despite these limitations, our integrative approach provides several key advances. First, we demonstrate that sophisticated analytical methods (NMF, MOFA) substantially outperform conventional approaches for patient stratification in complex diseases. Second, the convergence of multiple analytical pipelines on integrin signaling and lipid metabolism pathways provides robust evidence for their pathophysiological importance. Third, the identification of a multi-marker panel offers immediate translational potential for risk stratification.

## Conclusion

This study demonstrates the power of integrating multiple advanced bioinformatics approaches—Non-negative Matrix Factorization, Multi-Omics Factor Analysis, and machine learning—to identify molecular signatures associated with long-term pulmonary dysfunction in humidifier disinfectant-exposed adolescents. Through comprehensive multi-omics profiling and sophisticated analytical frameworks, we have established a robust multi-biomarker panel comprising IGHV-70, LysoPC (16:0), and hexadecyl ferulate, which achieved 81.46% classification accuracy in distinguishing patients with varying degrees of lung function impairment (Restrictive and Obstructive groups). Our findings reveal convergent molecular pathways underlying disease progression, with integrin signaling and lipid metabolism emerging as central mechanistic themes across multiple analytical pipelines. The identification of disease stage-dependent biomarker-clinical correlations, particularly the progressive strengthening of associations between CD36 and pulmonary function parameters, provides valuable insights into the dynamic nature of this environmental lung disease. These molecular signatures not only offer diagnostic potential but also illuminate potential therapeutic targets, particularly within the PPARA-regulated lipid metabolism network. The clinical implications of this work are substantial. The proposed multi-marker panel represents a promising tool for early risk stratification, enabling clinicians to identify high-risk individuals before the onset of severe respiratory symptoms. This proactive approach could fundamentally alter the clinical management paradigm, shifting from reactive treatment to preventive intervention. Moreover, the strong correlations between biomarkers and spirometric parameters suggest their utility for longitudinal monitoring of disease progression, potentially guiding personalized therapeutic decisions. In conclusion, this investigation represents a significant advancement in our understanding of humidifier disinfectant-associated lung injury, providing both mechanistic insights and practical tools for clinical application. By bridging molecular profiling with clinical phenotyping, we have established a foundation for precision medicine approaches in environmental lung disease, offering hope for improved outcomes in this vulnerable population as they transition into adulthood.

## Acknowledgements

We thank the humidifier disinfectants exposed to pediatric patients and their families participating in the study. This work was supported by a grant from the National Institute of Environmental Research (NIER), funded by the Ministry of Environment (ME) of the Republic of Korea (NIER-2023-04-02-160) and the Research Program funded Korea National Institute of Health (2008-E33030-00, 2009-E33033-00, 2011-E33021-00, 2012-E33012-00, 2013-E51003-00, 2014-E51004-00, 2014-E51004-01, 2014-E51004-02, 2017-E67002-00, 2017-E67002-01, 2017-E67002-02, 2020E670200, 2020E670201). We acknowledge the use of Claude Opus 4 solely for linguistic refinement and grammatical corrections in manuscript preparation. All scientific content, data analysis, and intellectual contributions presented herein were developed independently by the authors without the use of generative AI tools. Figure 1 was created with BioRender.com (accessed 1 July 2025).

## Supplement Figure Legends

**Supplement Figure 1. Limited discriminatory capacity of K-means clustering for clinical group stratification.**

(A) Optimization of cluster number for the initial timepoint dataset using the elbow method, with subsequent visualization of clustering results (*k* = 4) projected onto the first two principal components. The PCA representation reveals substantial overlap between clinical groups, indicating insufficient separation of distinct patient populations. (B) Parallel analysis performed on the final timepoint dataset, demonstrating comparable clustering patterns with *k* = 4. The persistent intermixing of clinical groups suggests inadequate capture of underlying biological heterogeneity. (C) Silhouette analysis validating the optimal cluster selection (*k* = 4) for both datasets. Average silhouette widths of 0.39 (initial) and 0.51 (final) indicate moderate to weak cluster cohesion and separation. (D) Hierarchical clustering dendrograms constructed using Ward’s linkage method applied to K-means cluster assignments. Branch coloring corresponds to cluster membership: Cluster 1(black), Cluster 2 (cyan), Cluster 3 (light green), and Cluster 4 (purple). Color-coding denotes distinct groups (Group 1: black, Group 2: cyan, Group 3: light green, Group 4: purple). The dendrogram structure further illustrated the absence of clear demarcation between predefined clinical groups in both initial (top) and final (bottom) datasets.

**Supplement Figure 2. Clinical correlation of biomarkers in other groups.**

The heatmap represents correlations between biomarkers and clinical variables for the normal and severe groups at the initial timepoint. No significant correlations were identified between biomarkers and pulmonary function parameters in the normal group. Conversely, numerous significant clinical correlations emerged in the severe group. APOA1, APOA2, PPBP, and PF4 demonstrated strong positive correlations with FVC. PPBP also exhibited a strong negative correlation with FEV_1_/FVC. Furthermore, LIMS1 (Uniprot: P48059, isoform 2) showed significant positive or negative correlations with three clinical parameters: FEV_1_/FVC, FEF_25-75%_ and FVC. The red box highlights the clinical correlation pattern of CD36, indicating disease stage-dependent correlation dynamics.

