## Supplementary figures and images for "Biomarker Discovery via Integrative Multi-omics for Children exposed to Humidifier Disinfectant"

### Supplement Figures 1 & 2

#### Slide 1
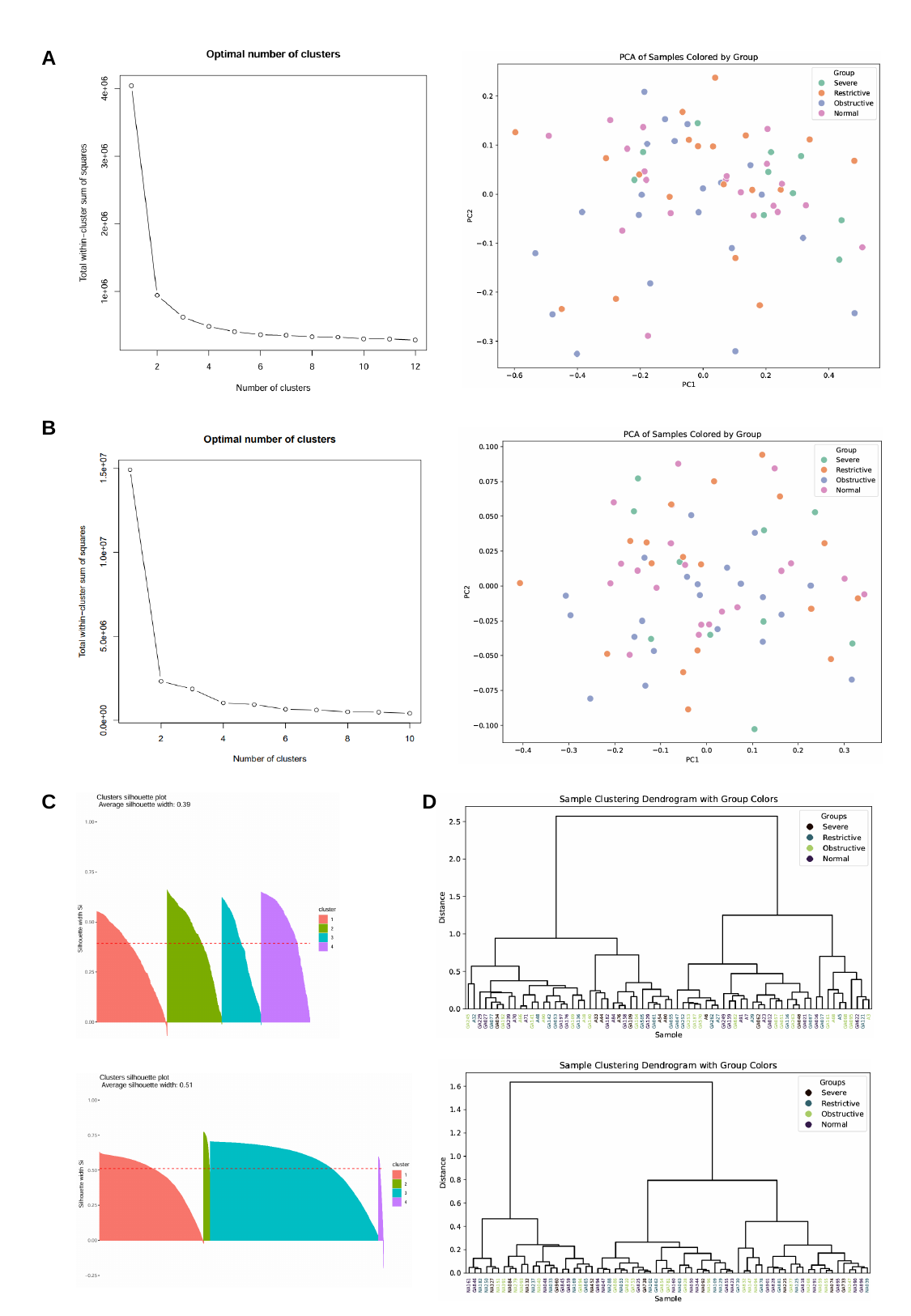

A
B
C
D

#### Slide 2
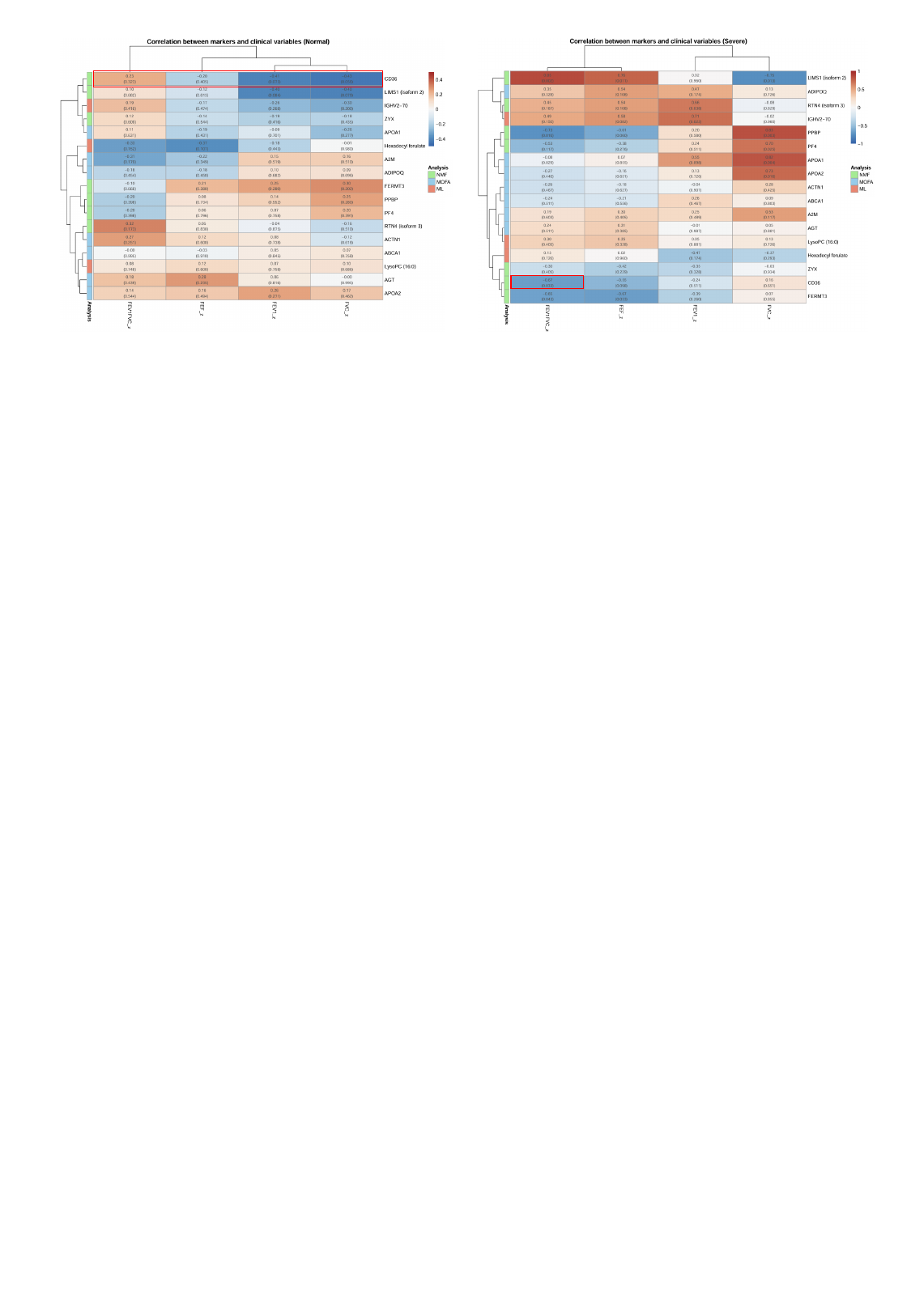
